# Inferring protein-induced spontaneous curvature of lipid bilayers from molecular simulation

**DOI:** 10.64898/2026.10.15.759481

**Authors:** Amirali Hossein, Melanie A. Brunet, Noah Englander, Jay Dadhania, Alexander J. Sodt

## Abstract

Proteins and lipids of a biological cell work together to reshape the membranes in which they reside. Membrane reshaping is necessary for vesicular traffic and maintenance of organelle shape and structure. *Curvature* is the fundamental mathematical description of the local shape of a surface. Through the well-established Helfrich/Canham/Evans model of bilayer shape, curvature sensing and induction are both consequences of a protein’s *spontaneous curvature*, the local curvature at which the Helfrich model energy is a minimum. This paper describes a straightforward approach for estimating the strength and sign of a bilayer-associated molecule’s curvature induction from molecular simulation: Correlating the molecule’s lateral position with curvature on thermally-fluctuating bilayer modes. Example demonstrations range from a large multi-pass coarse-grained transmembrane protein to an all-atom amphipathic helical segment. Quantification of these systems is placed in the context of experiments seeking the same strength and sign of curvature induction.

## Introduction

Cellular membranes are constantly reshaped by the cell as an essential component of homeostasis. Proteins are packaged into vesicles for trafficking to and from the plasma and organelle membranes as they are produced and recycled. Vesicle trafficking requires budding, fission, and fusion, all of which require dramatic deformations of the membrane. Membrane shape is quantified locally by *curvature*, denoted here as *J*, the mathematical measure of deviation from the planarity. The continuously defined curvature over a bilayer’s surface is its shape.

The propensity of lipids and proteins to generate curvature is a mechanical parameter intrinsic to the protein (1), lipid (2), lipid cluster (3, 4), leaflet (5) or asymmetric bilayer (6). It is typically called the *spontaneous* curvature, *J*_0_, a name most appropriate for leaflets and bilayers that can adopt curvature collectively, but also frequently applied to smaller elements based on their collective determination of leaflet spontaneous curvature.

The exact meaning of the term spontaneous (or intrinsic) curvature as used in literature related to membrane biophysics is somewhat ambiguous. It can refer to a total leaflet property determined from average spontaneous curvature of lipids that enters the Helfrich Hamiltonian as a constant determining the minimum of leaflet curvature energy. In this context “spontaneous” is the most appropriate as leaflets can spontaneously assume shapes, such as the inverse hexagonal phase (7). It also refers to a property of individual lipids or proteins quantifying that particular membrane element’s curvature preference, from which the leaflet spontaneous curvature is computed. In an additive model, the global properties of the membrane are assembled from the averages of individual lipid and protein properties. While the additive is a highly effective starting point, lipid spontaneous curvature is in part determined by the surrounding matrix, indicating non-additive coupling between components (8). Our working, generalized definition is that spontaneous curvature is a property of protein-lipid and lipid complexes that has a dependence on *q*, the wave number of membrane undulations.

Various mechanisms for generation and sensing of local curvature by membrane proteins have been identified. The protein or protein complex may imprint its shape on the underlying membrane, forming a *scaffold*. This mechanism is typically considered dominant for BAR domain–containing proteins, clathrin, dynamin, COPI, COPII, and ESCRT-III. Insertion of protein motifs such as amphipathic alpha helices can have a direct influence on membrane stresses via molecular mechanisms akin to a *wedge*, inducing either positive or negative curvature depending on insertion depth (9). At high coverage, bulky proteins can induce curvature by *crowding* (10).

The magnitude of coupling to curvature is a critical parameter for determining many membrane-mediated processes. Asymmetric distributions of spontaneous curvature of lipids alone can stabilize vesicle formation (11). Protein-induced mechanical changes can influence aggregation on the soft, fluctuating membrane (12–16). Clathrin-mediated endocytosis may involve many isolated molecular curvature mechanisms, from amphipathic wedging (9, 17), crowding, and scaffolding by the clathrin cage and its membrane-attached adaptors (18, 19). Extracting a clearly defined coupling constant that can be used regardless of molecular mechanism and lengthscale is critical for both model building and comparison to experiment. The following section justifies the choice of simulation observable (⟨*J*⟩*A*) as well as its equivalent model interpretation (*A*_p_*J*_0_).

### Simple linear-response curvature sensitivity from SPEX

**Sp**atial **Ex**tent (SPEX) methodology tracks a leaflet- or bilayer-bound molecule’s sampled curvature, ⟨*J*_*q*_⟩, on undulating modes with wavenumber *q*, to infer its spontaneous curvature, *J*_0_ from the linear coupling implied by the Helfrich model. The parameter *J*_0_ enters the Helfrich model of a region of the bilayer surface as

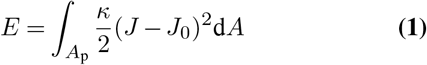

where for simplicity any influence of the protein on *κ* or 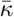 is not considered. Note that the sign of *J*_0_ is arbitrary to some degree, depending on the orientation of the protein in the bilayer and the sense targeted by the model. Here *J*_0_ is positive if the leaflet associated with the protein is convex with respect to headgroups — that is, positive if the protein sits on the outside of a vesicle. For a flat patch of bilayer (uniformly *J* = 0) the derivative of the patch energy with respect to curvature is

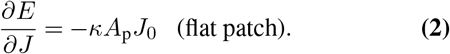

Leibler’s foundational theory that a protein’s curvature coupling to bilayer undulatory modes will reduce the apparent bending modulus (20) uses as its coupling coefficient

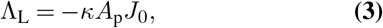

as shown in the Methods, this coupling yields for the curvature dynamically sampled by a protein ⟨*J*⟩ per mode *q*

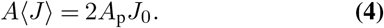

for a bilayer model (Eq. 1) or

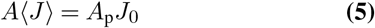

for a leaflet model. With the assumptions in Eq. 1 the result is independent of the material stiffness because the coupling coefficient and elastic coefficient are both proportional to *κ*; an increased protein bending modulus relative to the bulk would be equivalent to an increased *J*_0_ or *A*_p_. This paper describes critical technical details for extracting the linear coupling from a simulation, as well as results for a range of realistic test systems. Comparing SPEX results to experiment requires analysis of how curvature coupling has been reported in the literature.

### Experimental determination of curvature coupling, J_0_

Model predictions and experiments of curvature sensitivity can be compared quantitatively using the formalism here if the conditions are precisely controlled. Arguably the gold standard of experimental protein spontaneous curvature determination is to pull a narrow tube/tether (21) from a GUV (22) and quantify the enrichment of fluorescently tagged protein on the highly curved tube. The diffraction-limited tube radius is implied by the fluorescent magnitude of the tube relative to the flat GUV or from the force necessary to pull the tube, given the bending modulus (1). These pulled-tether experiments clearly show varying regimes between a dilute protein strongly enriching and coat formation on the tube at higher coverage (1). In Ref. (23), the composition-curvature coupling parameter Λ_B_ describing intensity enrichment 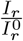:

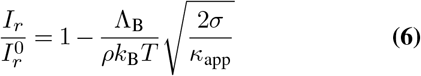

is measured to be -146 pN for the epsin N-terminal homology domain, a potential positive-curvature generator in clathrin-mediated endocytosis. Here the Baumgart lab follows a similar formalism of Ref. (20) in which the apparent bending modulus *κ*_app_ is derived for coupling constant Λ_L_ and coverage Φ with

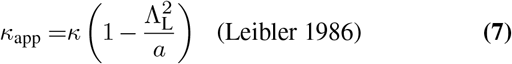

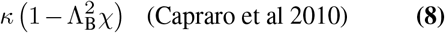

where here for Leibler’s softening expression

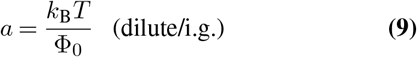

and Φ_0_ is the average density. Here Leibler allows for generality in protein-protein attraction with the constant *a* that modulates variations from the average density. For Capraro et al’s relation, surface coverage *χ* is determined in terms of the density of PIP2 lipids on the surface, rather than the ENTH footprint, with 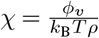, where *ϕ*_*v*_ is the PIP2 mole fraction and *ρ* is the lipid density 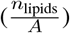. It follows that

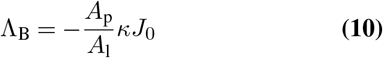

and has units energy/length (e.g., pN) where here the sign is fixed such that negative Λ_B_ implies a positive spontaneous curvature. If 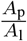 is (for example) three (that is, assuming ENTH has a footprint equivalent to three lipids), the spontaneous curvature implied by Λ_B_ = −146 pN is *J*_0_ = 0.59 nm^−1^ with *κ* = 20 *k*_B_*T* . This high positive curvature is consistent with the range of magnitudes predicted by the model of Campelo, McMahon, and Kozlov (9) (up to 0.75 nm^−1^) for amphipathic alpha helices similar to that of ENTH.

Relations between *J*_0_, Λ_L_, and Λ_B_ are summarized in Table 1.

**Table 1.** Equivalent expressions for curvature sensitivity.

| Bilayer/curvature mode coupling | $J_0$ equivalent | $\Lambda_L$ equivalent | Example Ref. |
| --- | --- | --- | --- |
| $\bar{C}_0$ or $J_0$ | - | $-\kappa A_p J_0$ | Sorre et al (1) |
| $\Lambda_B$ | $-\frac{A_l \Lambda_B}{A_p \kappa}$ | $A_l \Lambda_B$ | Capraro et al (23) |
| $\Lambda_L$ | $-\frac{\Lambda_L}{A_p \kappa}$ | - | Leibler (20) |
| $A\langle J \rangle$ | $\frac{A\langle J \rangle}{2A_p}$ | $-\frac{A\langle J \rangle \kappa}{2}$ | this work |

### Alternative simulation determination of J_0_

Figure 1 graphically summarizes available methodology for estimating the coupling of a protein to bilayer or leaflet curvature, including the SPEX method applying Eq. 4.

**Fig. 1.**
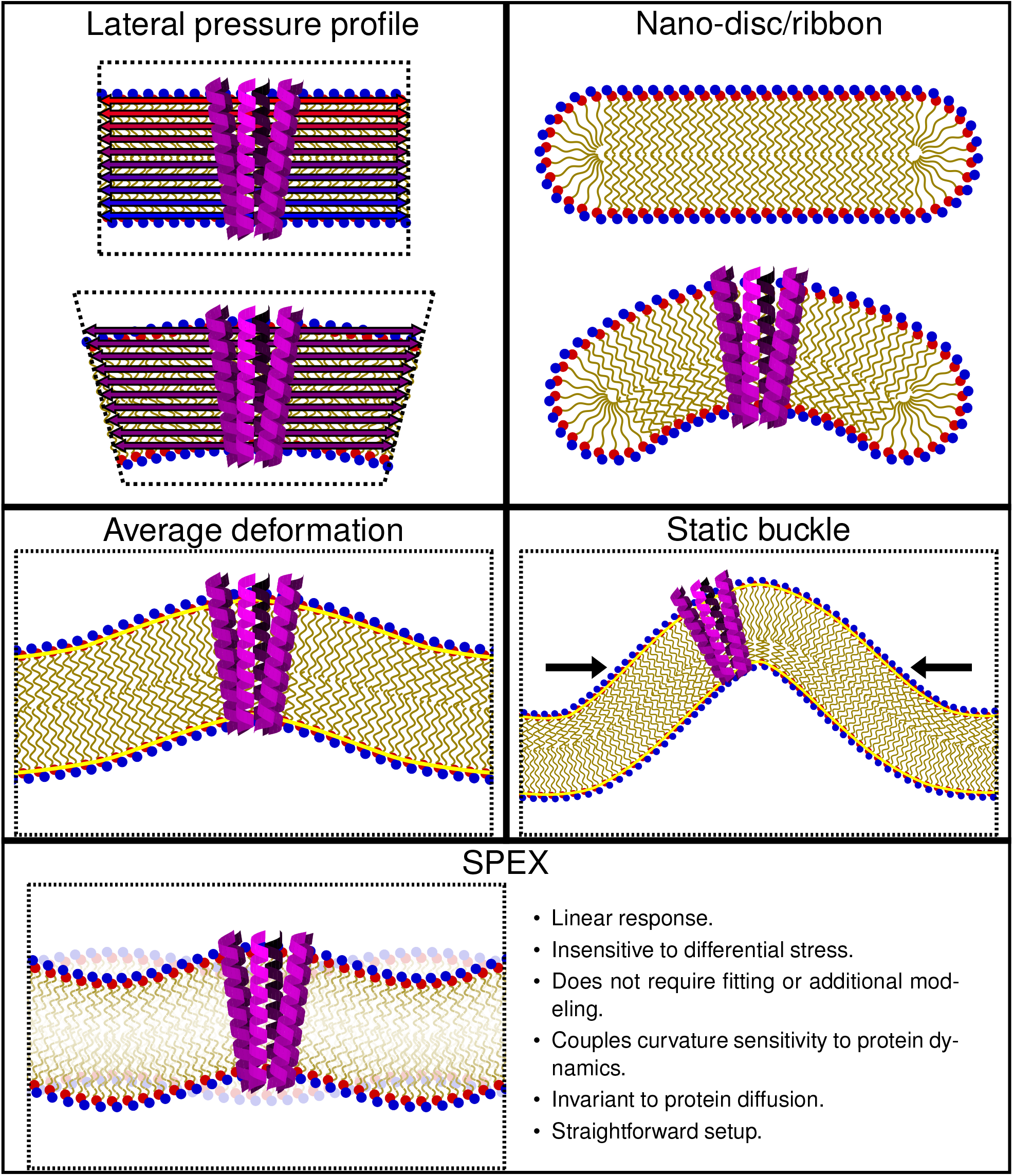
An overview of simulation methods for computing protein spontaneous curvature. **Lateral pressure profile** – With planar geometry enforced by periodic boundary conditions, proteins stress the surrounding bilayer, leading to static forces that can be translated into a curvature-free-energy derivative. **Nano-disc/ribbon** – with a free edge, a disc or ribbon is free to adopt high curvature induced by a protein. **Average deformation** – Proteins will induce a static deformation that is dependent on spontaneous curvature. **Static buckle** – Inducing a static buckle by applied tension creates high curvature to which a curvature-sensitive protein will localize. **SPEX** – This paper’s methodology. The protein couples to the local curvature of a fluctuation dependent on its spontaneous curvature.

#### The lateral pressure profile

The value of spontaneous curvature as a bilayer or leaflet property can be determined from the lateral pressure profile of the bilayer. Lateral pressure profiles indicate where there are contrasting lateral positive and negative tensions leading to curvature torques (related to derivatives of free energy, calculated using the Helfrich Hamiltonian, with respect to curvature).

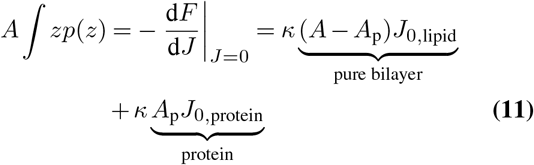

where *p*(*z*) is the lateral pressure profile. The change in *κAJ*_0_ attributed to an embedded protein or biomolecule is 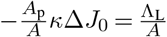, where Δ*J*_0_ is the spontaneous curvature difference between the protein and the matrix. In Refs. (24, 25), the lateral pressure profile of a bilayer was analyzed with and without an added amphipathic helical peptide. The change in the leaflet curvature free-energy derivative 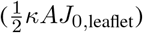 implied the spontaneous curvature of the added peptide, given the fraction of simulation containing the peptide 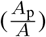. Equation 11 is derived from a virtual curvature deformation of the simulation unit cell, similarly to how the isotropic pressure is computed; this requires a well-behaved *contour* that determines how molecular forces are assigned spatially to *p*(*z*) (see e.g. Ref. (26)). A serious drawback is that the curvature derivative is sensitive to an imbalance of lipid counts between leaflets (27), although there are tools to aid in the choice (28). Moreover, implying *κ*_*m*_*AJ*_0,leaflet_ requires the assumptions of a model in which mechanical properties can be derived from the average of constituent properties.

#### Nano-discs and ribbons

Refs. (29–31) demonstrate the striking curvature induced by a protein on lipid nano-discs. A major advantage of this approach is the clear effect of the protein on a disc’s curvature, which, if constructed appropriately, can be taken well beyond the small deformation/linear response regime of a planar lipid deformation. A serious drawback is that, unlike the model energy of a sinusoidal planar deformation, the disc’s energy requires a sophisticated modeling approach that is subject to systematic errors, although edge-preferring lipids and specially designed asymmetric ribbons (32) are available to ease this modeling task.

#### Redistribution on a bilayer buckle

Refs. (33–35) apply static bilayer buckles induced by anisotropic tension to sample a protein’s curvature localization that is a direct result of *κA*_p_*J*_0_. The approach shares the advantage of the nano-disc approach that a large curvature is induced, leading to strong coupling. The energetic model of a buckle produces fewer ambiguities to modeling; indeed the predictable shape of a buckle can be used to extract the pivotal plane of bending (36). However, unlike the coupling between a single protein and undulations, quantifying redistribution on the buckle requires sampling the protein diffusion timescale.

#### Interpreting the average deformation

Refs. (37–39) interpret curvature features of the average deformation around a protein. Such an approach allows for fine details of the membrane deformation to be extracted, including the lipids near the protein boundary. While a continuum fit to the deformation is similarly difficult to the nano-disc above, with the H_min_ method the *q*^2^-weighted Fourier-transform of the average deformation is equivalent to the curvature sampled — and so these two approaches are closely related. The advantage of the SPEX approach is the ease of both computation and interpretation at longer wavelengths, that is, interpreting the total bilayer curvature deformations away from the protein.

Figure 2 shows the product of the SPEX curvature-analysis workflow. The area-weighted *spontaneous curvature spectrum A*_p_*J*_0,*q*_ is computed for a membrane-associated inclusion (here a lipid (40), a peptide, or a protein) from the correlation of an inclusion’s lateral position with local curvature, through Eqs. 4 or 5.

**Fig. 2.**
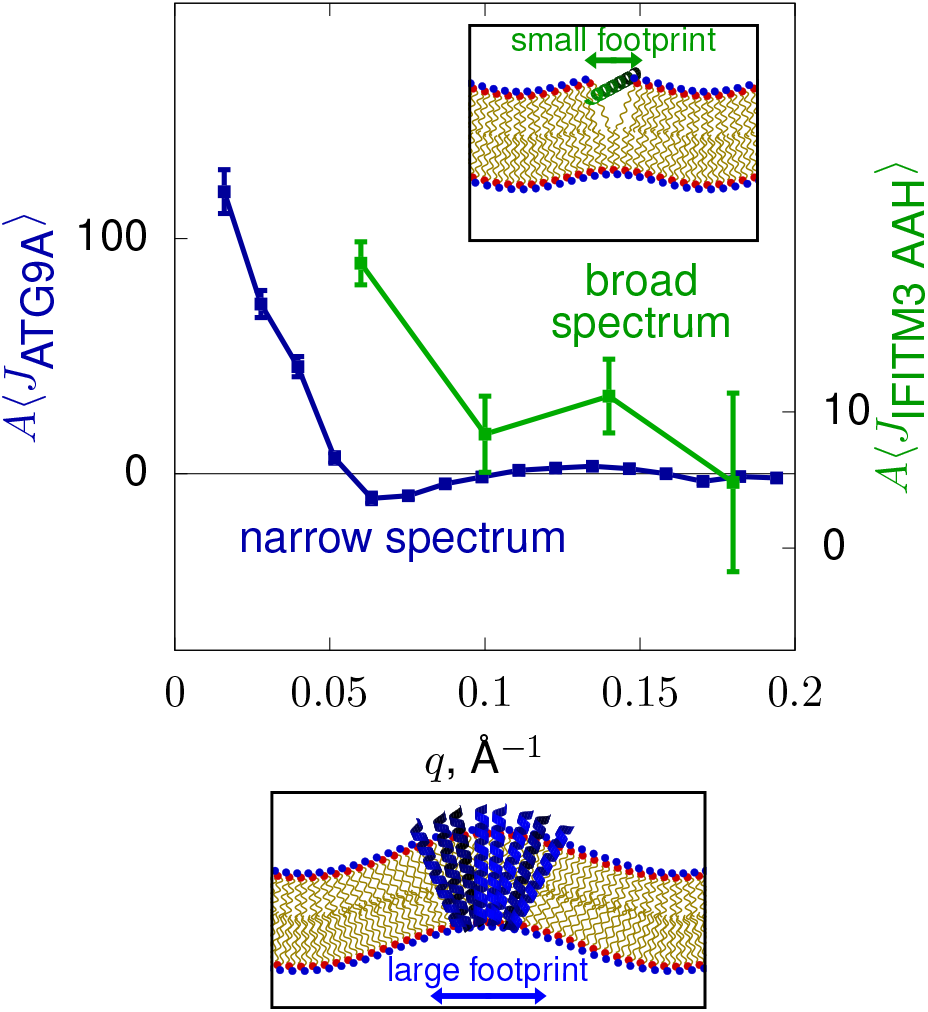
An overview of the primary result of the methodology: Length-scale (*q*) dependent spontaneous curvatures of a large multipass transmembrane protein critical for reshaping membranes (41) in autophagy (ATG9A) and a short, amphipathic segment of the antiviral immune system protein IFITM3 (IFITM3-AH). ATG9A induces curvature with a large spatial extent, leading to a narrow spectrum in Fourier-space. In comparison, IFITM3-AH, with its relatively small real-space footprint, has a broad spectrum in Fourier-space.

Demonstrations of the method here include

- A large multipass transmembrane protein from the autophagy machinery previously shown to induce qualitatively large curvature (ATG9A; coarse-grained Martini model).
- A peripherally attached C2 domain from synaptotagmin-1, a regulator of membrane fusion (all-atom model).
- An amphipathic alpha helical segment of an innate immune system factor that can be upregulated in response to viral infection (IFITM3; all-atom model).
- An idealized single-pass nearly symmetric transmembrane protein as a negative control (WALP (42); coarsegrained Martini model).

## Methods

### Protein spontaneous curvature from SPEX

The deformation of nearly planar upper (+) and lower (−) leaflets is expanded in a Fourier series, here with mode along *x* with integer *j*_*x*_:

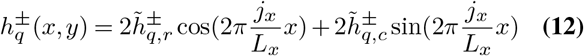

and for the bilayer midplane

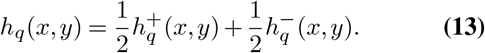

The base Helfrich model energy for a planar configuration is computed as

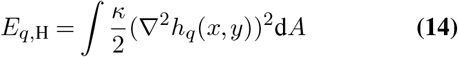

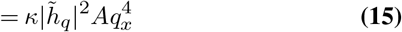

With 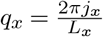. Here *E*_*q*,H_ is the energy for two independent degrees of freedom (the real and complex parts of 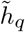).

The Helfrich model is then perturbed by the introduction of a protein at *x*_p_, *y*_p_, either as a *bilayer* perturbation

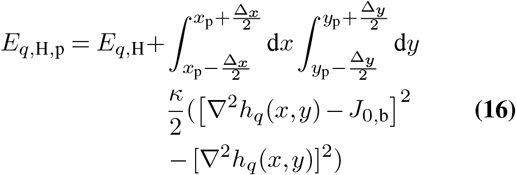

or (upper) *leaflet* perturbation:

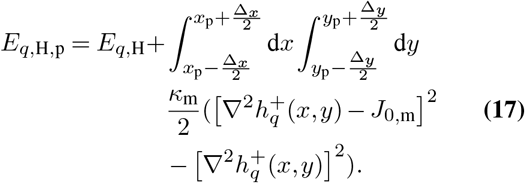

where here 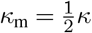. Simulating a single protein on a bilayer and sampling the curvature at the location of the protein yields ⟨*J*⟩. The expectation from the above model is computed through the standard statistical mechanical approach as

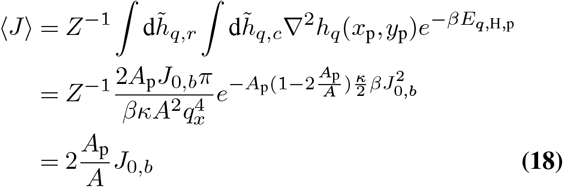

with

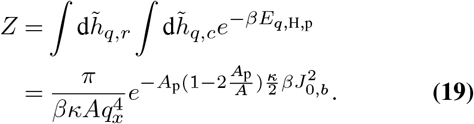

The curvature sampled for the upper leaflet model (Eq. 17) is halved, 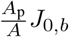, equivalent to a protein with half the projected area, or a lipid in a single leaflet (40).

### Interpreting the sampled curvature ⟨*J*⟩

With a model of *A*_p_ the spontaneous curvature can be inferred from ⟨*J*⟩ using Eq. 4 as 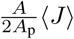 for a bilayer model of protein influence or 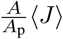 for a leaflet model. Alternatively, the ambiguity of *A*_p_ can be avoided entirely by considering a bilayer *patch*-centered view of spontaneous curvature. For simplicity, consider a bilayer of symmetric composition such that spontaneous curvature is zero. A protein then samples curvature ⟨*J*_*q*_⟩ on a periodically-replicated cell (patch) of projected area *A*. The spontaneous curvature of the patch is then a sum of the background (zero) and the protein spontaneous curvature (*J*_0_) multiplied by its simulated area weight 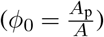:

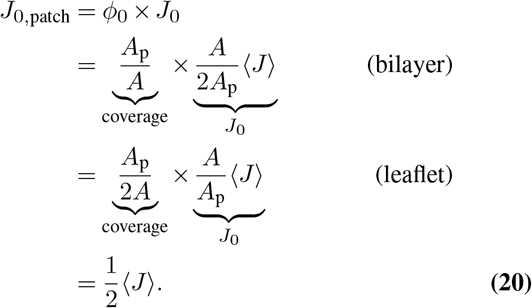

This view removes the ambiguity about the bilayer or leaflet influence of the protein, as well as its projected area. Arbitrary concentrations (*n*^′^ proteins yielding coverage *ϕ*^′^) and patch sizes (*A*^′^) can then be considered as:

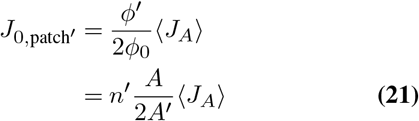

where here primed variables indicate the target conditions, and the subscript *A* in *J*_*A*_ indicates the protein was sampled on area *A* in the source simulation from which the target conditions will be derived. A straightforward presentation of protein spontaneous curvature is then to report *A* ⟨*J*⟩. With this value, patch spontaneous curvature is computed by Eq. 21. Enrichment of the protein as a function of curvature, relative to a flat region, is computed as

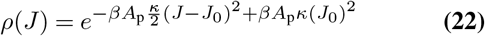

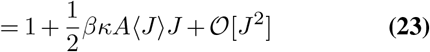

where the model has been defined for a bilayer (Eq. 22), Eq. 23 is equivalent for a bilayer or leaflet model. Note that for both enrichment and spontaneous curvature induction, the *q*-range over which ⟨*J*⟩ is averaged should be consistent with the lengthscale of the question. That is, for regions in which curvature varies rapidly, high *q* may be relevant.

### Interpolating membrane shape by minimizing the curvature energy, the H_min_ method

This section describes the method for sampling a protein’s curvature using the `MembraneAnalysis.jl` software (43).

Undulations of flat membranes in simulations are typically analyzed by discretizing the surface into a grid and assigning values for quantities such as surface height to each cell by averaging values from molecules positioned within the cell. The conventional method for dealing with cells without any lipids inside is to assign the average value over adjacent cells to that cell. This method is practical if the molecular density and grid size relate such that instances of empty cells are rare. There will be many contiguous empty cells for a protein spanning multiple discretization cells. The H_min_ method assigns the height for empty cells as free parameters and minimizes the Helfrich energy.

Figure 3 shows a snapshot from a coarse-grained simulation of a single ATG9A protein in a POPC membrane, with simulation data taken from Ref. (41) where the model was demonstrated to induce strong curvature. In this figure the blue circles show the position of PO4 beads of the Martini lipids in the upper leaflet and the grid is based on ≈15 Å cell side length. The cells with no lipids are orange and the cells in yellow are within *r*_mask_ = 20 Å of the backbone (BB) atoms of the protein. Here *r*_mask_ is the range in which the cell is *masked*, either because the cell is empty, or because the lipids are suspected to adapt to the shape of the protein and are thus likely inconsistent with an elasticity model.

**Fig. 3.**
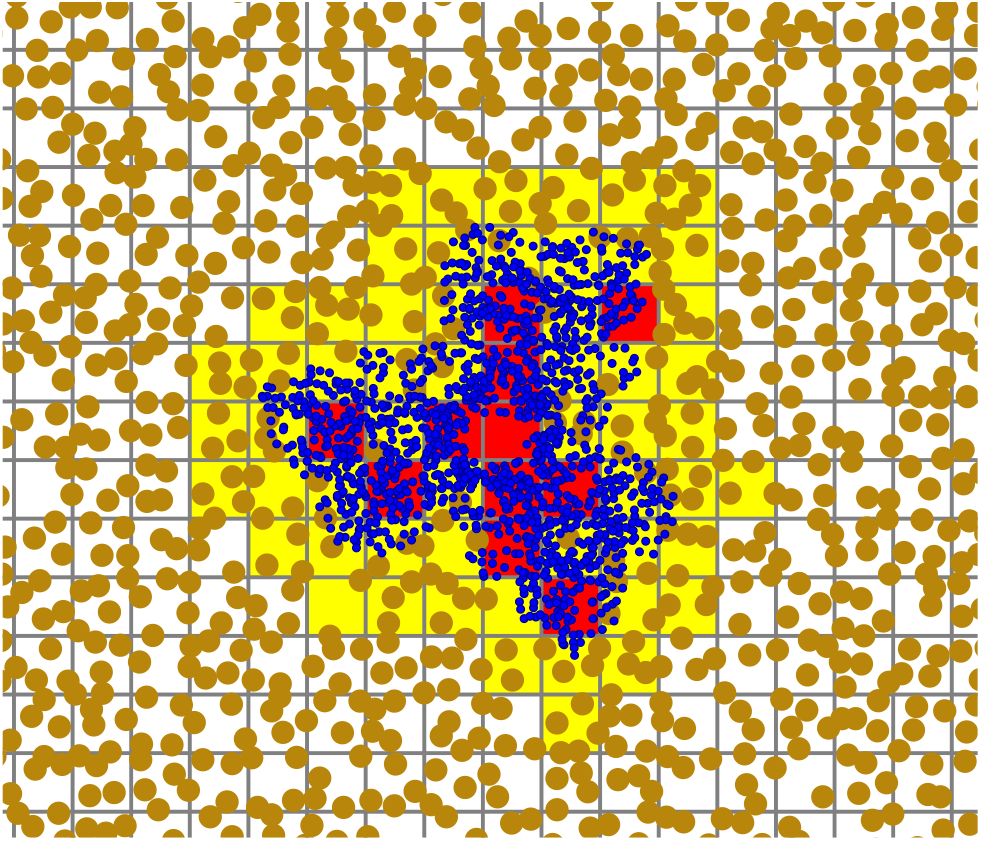
A snapshot from a simulation (41) of ATG9A in a POPC membrane, representing ca. 25% of the total simulation area. Grid spacing is 15 Å, the size of points used for Fourier analysis.

The surface height matrix 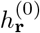 is constructed with the average height of the reference lipid atoms in each cell. The set of cell index pairs (*i, j*) corresponding to the excluded cells is *M* . The *M* cells of 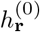 are then set to zero.

We calculate 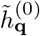, the discrete Fourier transform of this matrix. Next, for each of the excluded cells *m* we calculate the trivial discrete Fourier transform of a matrix of the same size as 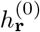 with all elements zero except the excluded cell, 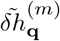. From linearity of the Fourier transform we can write:

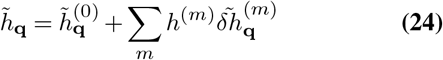

where *h*^(*m*)^ is the unknown height of the *m*th excluded cell. Setting to zero the derivative of the model energy *E* with respect to all unknown heights simultaneously yields *m* equations of the form:

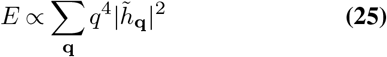

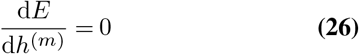

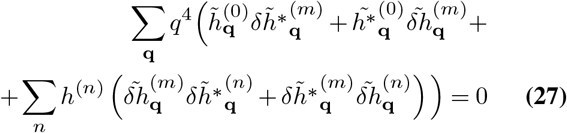

If *Z* is a vector of the height values of the excluded indices (*Z*_*m*_ = *h*^(*m*)^), we can write this system of equations as the following matrix equation:

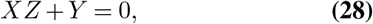

where *X*_*m,n*_ is

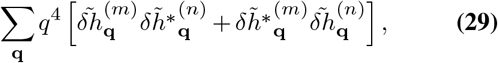

and *Y*_*m*_ is

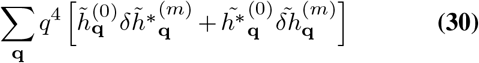

We can recover the energy-minimized height values of the excluded cells by inverting *X* from

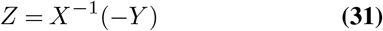

with *X*^−1^ computed using the Moore-Penrose pseudo-inverse such that, in the case of linear dependence, no additional height is added.

`MembraneAnalysis.jl` identifies the excluded cells and calculates height values separately for each leaflet. Heights in excluded cells are filled based on the H_min_ method. The reconstructed leaflet height matrices then determine the thickness and midplane height fluctuation modes from the mean (midplane height) and difference (thickness) of the two.

### Simulations

#### ATG9A

Simulations of the ATG9A trimer were published in Ref. (41) and provided by the authors for re-analysis. Briefly, a single ATG9A trimer was simulated in a POPC membrane with dimensions of approximately 500 × 500 Å with GRO-MACS 4.5.5 and the coarse-grained (CG) MARTINI 2.1 forcefield at constant semi-isotropically applied pressure (1atm) and constant temperature (298K) with the Berendsen barostat and thermostat, respectively. Non-bonded interactions were calculated using shifted Lennard-Jones Coulombic potential, both cut off at 12 Å, with a relative dielectric constant of 15. The duration of the trajectory was 1*µs*. Standard errors are computed by dividing the simulation into three blocks.

#### IFITM3-AH

Simulations of Wild-type (WT) IFITM3 amphipathic alpha-helical (AH: amphipathic helix) peptides were previously published in Ref. (44), which included current author AJS. The simulations were re-analyzed in the current work to compute their curvature induction. Amphipathic segments of the full protein were built as ideal alpha-helices. Two peptides (one on each leaflet) were placed near the bilayer-water interface. The bilayer was composed of 70:30 mol% POPC:cholesterol with 200 lipids per leaflet, and the solution was composed of 50 waters per lipid with 150 mM KCl. The system was minimized and briefly equilibrated using NAMD with the standard CHARMM-GUI protocol, then converted to Amber format using ParmEd. The simulations used the CHARMM36m all-atom force field, and Amber20 version of pmemd built for graphical processing units. Constant pressure of 1bar was maintained by a Monte Carlo barostat. Constant temperature was maintained at 310.15 K using Langevin dynamics with a 1ps_−1_ damping coefficient. Non-bonded forces were switched off between 10 and 12 Å. Covalent bonds involving hydrogen were constrained using the SHAKE and SETTLE algorithms. Long-range electrostatics were calculated by particle mesh Ewald summation. The system was simulated for 1 *µ*s. Standard errors are computed from averages over replicas.

#### Syt-1

Systems containing a single copy of each Syt-1 C2 domain (human gene) starting from the AlphaFold structure in a POPC:POPS (75:25 mol%) bilayer were built using CHARMM-GUI Membrane Builder tool with the CHARMM C36m all-atom force field. The systems were minimized and briefly equilibrated using NAMD with the standard CHARMM-GUI protocol, then converted to Amber format using ParmEd. The C2A domain was run for 1.5 microseconds; C2B was extended to 7.5 microseconds.

The production runs were maintained at constant pressure (of 1 bar) and temperature (of 310.15 K) using the Monte Carlo barostat and Langevin dynamics (with 1 ps_−1_ damping coefficient), respectively. Non-bonded forces were switched off between 10–12 Å, long-range electrostatics were calculated by PME, and covalent bonds involving hydrogen were constrained with the SHAKE and SETTLE algorithms. Standard error was computed from 300 ns blocks for C2A, 2 microseconds blocks for C2B.

#### WALP

The WALP sequence (42) was built as GWW(LA)_8_LWAA using the CHARMM-GUI Martini Maker module, starting from an initial ideal alpha helix oriented along *z*. The system contained 300 POPC lipids per leaflet, sufficient to sample curvature induction at relatively long wavelengths. Dynamics were computed using Gromacs, consistent with the Martini 3 forcefield used here. Parameters were taken from CHARMM-GUI consistent with recommendations by the forcefield: 20 femtosecond timestep, temperature controlled using the v-rescale algorithm, zero lateral tension controlled using a Parrinello-Rahman barostat (*τ*_*p*_ = 12.0 ps, compressibility = 3 × 10_−4_ bar^−1^, 1 bar *z* pressure). Three replicates were run for 10 microseconds for both neutral and charged termini. Standard errors are computed from averages over replicas.

## Results

### A large transmembrane protein: the autophagy protein ATG9A

Figure 4 shows the *q*-dependent spontaneous curvature of a coarse-grained (Martini) model of ATG9A. At the lowest *q* value sampled 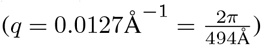, well below the size of the protein, area-weighted sampled curvature is 113 ± 7 Å. For a protein area *A*_p_ of ca. 6300 Å^2^, this corresponds to a (bilayer) spontaneous curvature of 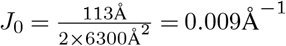.

**Fig. 4.**
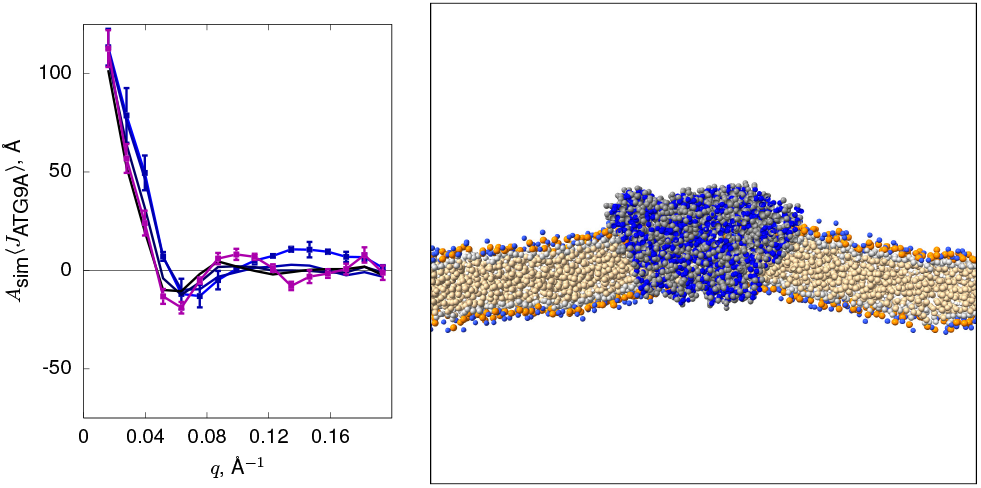
Curvature spectrum of ATG9A in a POPC membrane for *r*_mask_ values of 0, 10 Å, 20 Å, and 30 Å using the H_min_ method (ranging from bright blue to black), in addition to curvature spectrum calculated from the average deformation after applying H_min_ (purple). For clarity error bars are only shown for *r*_mask_ = 0; error bars for large *r*_mask_ are similar.

### Peripherally-bound C2 domains: Syt-1 C2A and C2B

Figure 5 shows the *q*-dependent spontaneous curvature of independently-simulated all-atom models of isolated C2A and C2B domains of SYT1. These domains are peripherally attached to the leaflet. Membrane binding is determined by insertion of a calcium binding loop as well as electrostatic contacts between anionic lipids and basic patches on the surface of the domain. Negative curvature is consistent with the argument proposed in Ref. (45), that the domain imposes its convex shape on the leaflet with its lipid contacts and embedded hydrophobic loop, leading to negative (concave with respect to head groups) leaflet curvature. There is a strong drop off in the curvature induced at higher 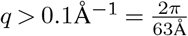 as *r*_mask_ is increased, consistent with the approximate size of the C2 domain (ca. 30 Åin length). We propose that this reflects the loss of information of these deformed anionic lipids in contact with the domain. At lower *q*, curvature is negative even with the contacting lipids masked (i.e., higher mask radius, dark curves in Fig. 5), indicating unambiguous curvature induction.

**Fig. 5.**
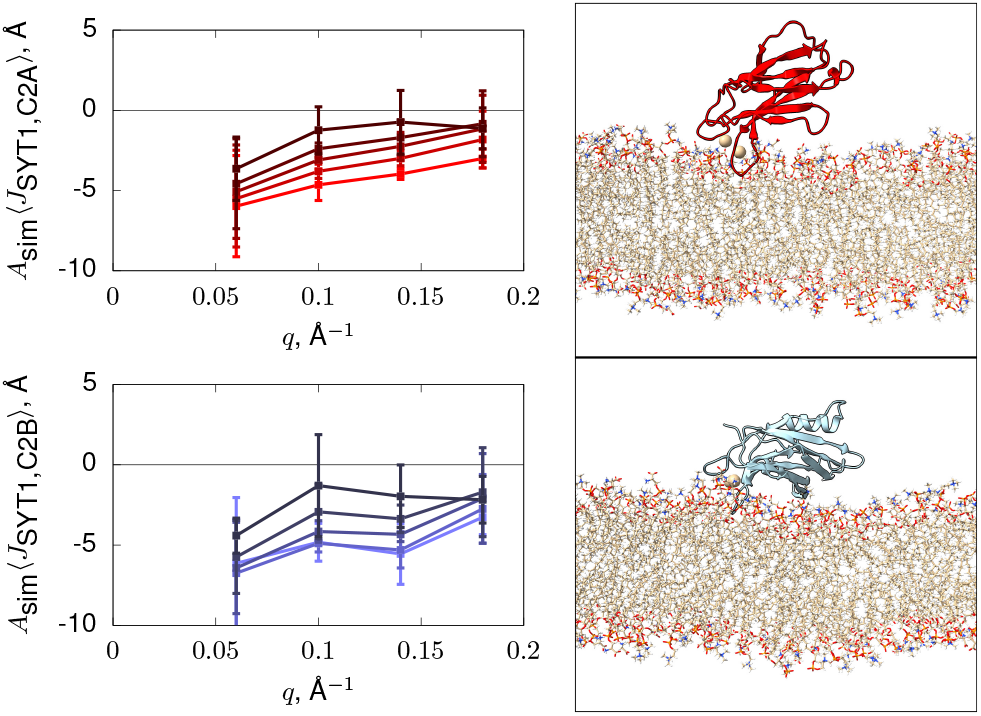
Curvature spectrum of SYT1 C2A (in red) and C2B (in blue) in a %75POPC:%25POPS membrane for *r*_mask_ values of 0, 5 Å, 10 Å, and 15 Å using the H_min_ method, with the brighter line representing smaller *r*_mask_ value.

### A single-leaflet adsorbed amphipathic helix: IFITM3-AH

Figure 6 shows the *q*-dependent spontaneous curvature of an all-atom model of IFITM3 amphipathic helix (IFITM3-AH), indicating positive curvature (as expected for an AH, but inconsistent with the interpretation of Ref. (46)).

**Fig. 6.**
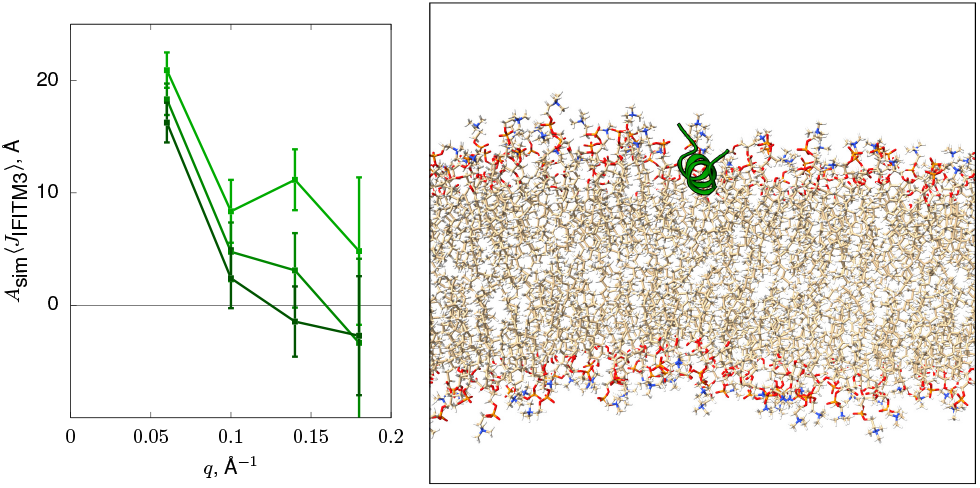
Curvature spectrum of IFITM3-AH in a membrane consisting of POPC and 30% cholesterol for *r*_mask_ values of 0, 5 Å, and 10 Å with the brighter line representing smaller *r*_mask_ value.

### A nearly symmetric single pass transmembrane helix: WALP

Figure 7 shows the *q*-dependent spontaneous curvature of a coarse-grained MARTINI model of WALP, indicating nearly zero curvature as expected from a nearly symmetric system.

**Fig. 7.**
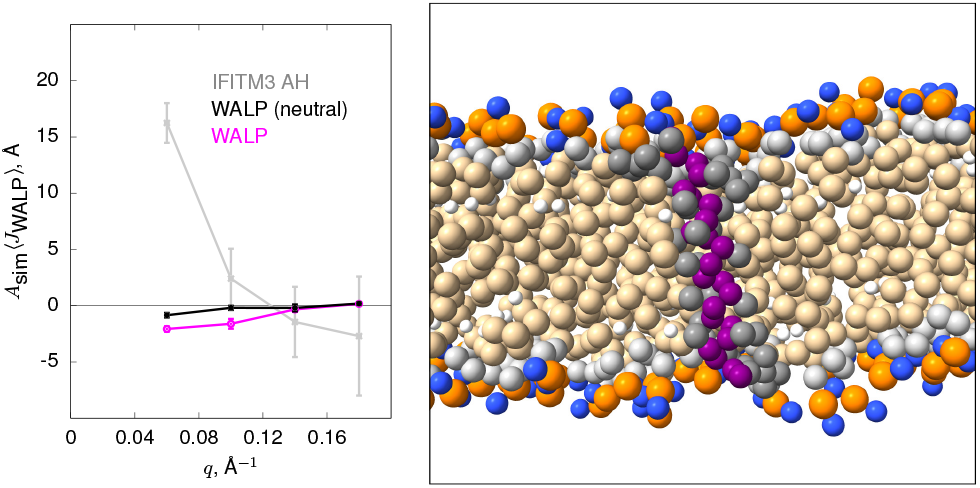
Curvature spectrum of WALP (both with charged and neutral termini) in a POPC bilayer with*r*_mask_ = 10 Å. The IFITM3 AH is shown for reference of magnitude.

### Validation computing the bending modulus with information partly removed

Figure 8 shows a validation of the procedure for the challenging estimation of the bending modulus, which is typically sensitive to non-elasticity artifacts in the fluctuation spectrum (e.g., from lipid tilt (47)). Three replicas of a 200 lipids per leaflet bilayer of CHARMM C36 all-atom POPC were simulated for one microsecond per replica. For each frame, a random subset of cells were marked to be excluded from the calculation of the fluctuation spectrum for a range of fraction of excluded cells (the abscissa/*x* axis of Fig. 8). Briefly, the bending modulus was computed using the SPEX bending modulus methodology (48), in which the apparent sampled curvature of lipid’s atoms is plotted against the height (*z*) of those atoms in a planar reference state. Softer bilayers have greater undulations and the method computes the *z*-dependent bias of a lipid to be oriented away from positive curvature, which is proportional to the undulation. Ref. (48) has a complete description of the approach to extract the bending modulus. In addition to the random exclusion of information, a circular selection of cells was also tested, providing a comparison similar to that of the footprint patch of a multi-pass transmembrane protein (like ATG9A). For the H_min_ method, the bending modulus extracted depends weakly on the fraction of cells below 50%, consistent with the relatively low *q*-range of modes used for the analysis (*q <* 0.08 Å^−1^. The discrepancy is markedly higher for the circular exclusion, presumably because the grouped cells lose important information for the low *q* undulations in the analysis. Nevertheless, the error is small (+4%) for a circular patch that excludes quite a large fraction (20%). Note that the error is systematically toward stiffer bilayers for both interpolation and H_min_, as the interpolated height is an energy minimum so likely misses some information of both thermal fluctuations as well as the systematic perturbation of the protein. That is, the tendency will be to *underestimate* the coupling of a protein to curvature.

**Fig. 8.**
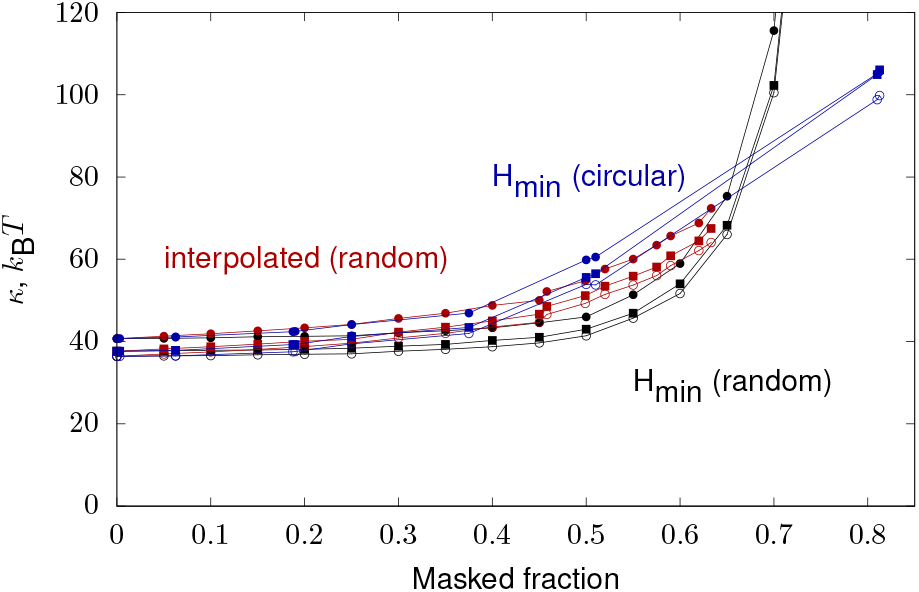
Validation of the H_min_ method for computing the POPC bending modulus by artificially excluding information from cells either randomly (black) or in a circular pattern (blue). The H_min_ method is compared with simple interpolation from populated neighbors (red). The H_min_ method has the desired flat behavior for low fractions but diverges significantly above 50% masked fraction. Interpolation is adequate for handling small numbers of unpopulated cells. The H_min_ method is effective even for large numbers of unpopulated cells in a circular pattern, which cannot be trivially interpolated from neighbors.

## Discussion

### Protein spontaneous curvature spectrum indicates protein’s local influence

Figures 4, 5, 6, and 7 show the curvature sensitivity for four systems spanning a large multi-pass protein to a single-pass protein with no expected curvature induction. In each case the *q*-dependence of the curvature is consistent with protein shape and size (see Fig. 2), in which large proteins have narrow spontaneous curvature spectra (Fig. 4) while small peptides (Fig. 6) have broad spectra, indicating that curvature is induced even at small length-scales.

Peripherally-bound Syt-1 C2 domains have flat spectrum suggesting influence at short lengthscales, although curvature induction is weaker for the system across *q* (see Fig. 5). Previous work using nano-ribbons indicated the C2 domain induced positive curvature, likely by a wedge mechanism (49). In the present simulations, the helix did not insert in an orientation to induce curvature; instead, negative curvature appears to be induced by protein-lipid contacts with the convex C2 domain, similarly to the model proposed in Ref. (45).

The 14 residue amphipathic helix of IFITM3 is critical for the antiviral activity of full length IFITM3 (50). Experiments suggest it induces negative curvature in giant vesicles (46). Here Fig. 6 shows that the purely alpha-helical model segment induces positive curvature as expected by the wedge mechanism, and consistent with results derived from lateral pressure profiles (24) of amphipathic helices. In the all-atom model, the sequence is insufficiently hydrophobic to embed deeply and induce negative curvature (9, 24, 34, 51). Discrepancies between the experiment and simulation may result from difference in conditions: the simulations are artificially precisely-controlled, reporting on isolated peptides embedded in the bilayer leaflet as complete alpha helices.

### Negative control

The single pass artificial WALP induces negligible curvature, consistent with its nearly symmetric structure. When made even less polar by neutralizing the N- and C-termini, the already low curvature sensitivity further diminishes (Fig. 7).

### The H_min_ method recovers curvature generation even with limited lipidic coverage

Spontaneous curvature spectra computed using increasing values of *r*_mask_ for ATG9A, synaptotagmin-1 C2 domains, and the AH helix indicate that curvature induced at low *q* is insensitive to the mask radius, while at high *q* implied spontaneous curvature drops dramatically in the case of ATG9A and slowly in the case of the IFITM3-AH. Recovering curvature at low *q* is consistent with the method retaining the full undulation shape at long wavelengths. Given the slope of curvature induction for ATG9A at low *q*, the patch size is not sufficient to display the full, zero-*q* curvature induction. A linear extrapolation of the lowest three *q* bins yields an intercept of 156 ± 3 Å, 38% larger than the q=0.016 Å^−1^ point. For the IFITM3-AH, the *q* = 0 intercept is 40 ± 6 Å, nearly twice as big as the *q* = 0.06 point. Minimal curvature induction at higher *q* (see, Fig. 4 above *q* = 0.05 Å^−1^ and Fig. 6 above *q* = 0.1 Å^−1^) is consistent with the area footprint of these peptide/protein systems.

In protein-free systems, randomly ignoring lipids when computing the bending modulus only increased the observed stiffness slightly (Fig. 8). Indeed, even for a large circular patch of removed information (20% of the area) the bending modulus matched closely (+4%).

### The SPEX method quantifies a simple model of curvature induction and enrichment

Eq. 23 evaluates the Boltzmann factor at linear order, consistent with the linear coupling determined from the model. The ATG9A curvature induction (113 Å) indicates it will enrich to +50% at a curvature of (250 nm)_−1_, that is, a 500 nm diameter tube or a 1 *µ*m diameter vesicle (with *κ* = 20*k*_B_*T*). Implying spontaneous curvature of the *simulated* patch from sampled curvature 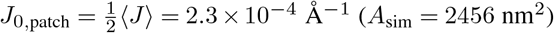, the spontaneous curvature of a target patch is inferred from Eq. 21. For example, if there are 25 copies of ATG9A (52) on a 45 nm transport vesicle 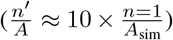, along with other curvature inducing proteins (e.g., BAR), the spontaneous curvature is (45 nm)^−1^, that is, comparable to the strength required to induce the formation of a small transport vesicle. Indeed, following Leibler’s instability theory, the concentration required for protein-curvature localization to collapse on a shared preferred curvature is

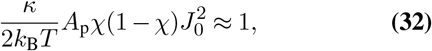

that is, the condition for the bending modulus to be zero (Eq. 7). In the linear regime (neglecting 1 − *χ*, the contribution of the protein itself to background spontaneous curvature) the formula simplifies to

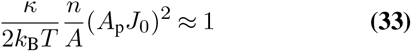

where 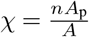. Inserting 2*A*_p_*J*_0_ = 113 Å, the coverage is higher than would be necessary to induce a curvature instability from ATG9A alone. Given that the coverage from ATG9A in the source organelle of the transport vesicle is lower, this does not strongly indicate ATG9A as a coatomer sufficient to form the vesicle; however, it does suggest that addition of the ATG9A vesicle pushes the composition of the targeted vesicle-absorbing organelle toward an instability that would indicate a major change in shape to accommodate ATG9A spontaneous curvature. Note that this analysis does not consider any interactions between proteins that are necessary to form a complete model of curvature-mediated aggregation (16).

IFITM3 and Syt-1 have far more modest area footprints and correspondingly less curvature induction. Localization of individual proteins is thus strongly reduced, with the curvature required to reach 50% enrichment proportional to *A*_sim_ ⟨*J*⟩. Where ATG9A enriches to 50% at 500 nm^−1^, with *A*_sim_ ⟨*J*⟩ nearly ten times less, IFITM3 requires ten times the curvature, e.g., a 200 nm diameter vesicle. However, an instability may be reduced at similar *χ*, which depends on *A*_p_. That is, a ten times lower *A*_p_ of IFITM3 would compensate for its ten times lower per-molecule curvature induction, relative to ATG9A. While the breakdown of 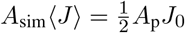 into *A*_p_ and *J*_0_ contributions does not emerge naturally from the model, a rough estimate of *A*_p_ can certainly be obtained from inspect of, for example, the occupancy of grid cells in Fig. 3.

## Conclusions

The paper demonstrates a method quantifying the linear-response curvature induction of four protein/peptide systems in simple lipid bilayers. Curvature sensitivity is implied by the sampling of a single protein/peptide on the thermal undulations of the bilayer, where sampling is proportional to spontaneous curvature and the area projection of the protein.

Large proteins (here, ATG9A) couple only to long wavelength modes to which they appear “point-like”, whereas smaller peptides can couple to shorter wavelengths (higher *q*). The linear coupling strength informs corresponding linear estimates of enrichment and curvature induction at target coverage and average curvature.

Given the relative ease of the simulation workflow and its statistical power, the method forms a natural complement to other techniques (reviewed above) that extract curvature coupling beyond the linear regime. The power of methods that extract curvature induction from, e.g., nano-discs and ribbons would be both more powerful and more interpretable if they also report where curvature induction differs significantly from the linear response estimate.

## Data sharing

Trajectories and structure files for IFITM3, Syt-1, and WALP simulations are available via Zenodo, at DOI 10.5281/zenodo.23345543.

## Acknowledgments

We are grateful for helpful discussions on curvature generation by ATG9A with Juan Bonifacino. Simulations of ATG9A were kindly provided by the authors of Ref. (41).

